# Application of Computer Vision Tools to Maize Genomic Data for Trait Prediction and Gene Discovery

**DOI:** 10.64898/2026.05.21.726890

**Authors:** Steven A. Higgins, Emily Anible, Mughilan Muthupari, Christopher Dibble, Robert W. Murdoch

**Author notes:** Correspondence: Robert W. Murdoch, Battelle Memorial Institute, 505 King Avenue, Columbus, OH 43201, USA.

## Abstract

Artificial intelligence and machine learning for computer vision (CV) and image recognition is a rapidly evolving field with multiple potential applications in plant genomics. While CV has been widely adopted by the research community for plant phenotyping and disease surveillance, applications of CV tools to plant genome analysis are underrepresented. CV tools may complement traditional statistical classification tools used in plant genomics, since CV perceives problems holistically rather than granularly (in terms of pattern recognition), which is particularly applicable to analysis of large, complex eukaryotic genomes. In this study, we report on a new strategy to apply existing CV tools to classify plant genotypes and predict genotype-phenotype relationships. A technique was developed for converting maize genome resequencing data into a set of images reminiscent of a quick response (QR) code. Several hundred maize genomes were processed and it was demonstrated that CV models can successfully categorize genome images into heterotic groups (accuracy and recall > 0.8). Models for classifying genome images into phenotypic trait groups (such as short, medium, and high plant height) performed with moderate success for the most heritable trait analyzed (ear height; accuracy and recall > 0.5). Querying model results permitted identification of genome regions that were important for model classification predictions. The CV model results revealed enriched metabolic pathways consistent with traits under consideration. Overall, our initial application of CV tools to plant genome analysis highlights its applicability to genomic data. Design of new CV architectures optimized for genome-derived images may further improve upon our initial results generated using only off-the-shelf CV tools optimized for unrelated image analysis tasks.

**Core ideas:**

- AI/ML computer vision (CV) tools were applied to encoded maize genomes
- CV image classification tools were able to successfully classify encoded genomes into heterotic groups
- Trait values of maize strain ear height could be predicted with moderate success
- Genome regions encoding plausible metabolic pathways used by the classifier were identified
- Recommendations for improved success of CV for genotype-to-phenotype are discussed

## INTRODUCTION

Genome-wide association studies (GWAS) are the de facto method for identifying genetic markers underlying plant phenotypes. Intense development of GWAS over the last two decades has seen incorporation of approaches to account for population structure and improve statistical calculation efficiencies, leading to ever-increasing reported links between genetic variants and traits (Yoosefzadeh-Najafabadi *et al*., 2022). In crops, functional alleles identified by GWAS have been successfully used to inform breeding programs to improve yields and disease resistances (Sun *et al*., 2020). Popular GWAS tools tend to remain firmly rooted in traditional statistical testing and modeling, essentially performing one-by-one tests between variants and traits (Shikha *et al*., 2021). The ever-decreasing costs of large genome resequencing and improved reference genomes allow for simultaneous evaluation of enormous numbers of variants. The statistical basis and large number of variants per study leads to challenges in establishing appropriate multiple testing corrections for p-value significance. GWAS also struggles to identify variants with weak effects or suites of variants that work in concert to drive phenotype (Tam *et al*., 2019, Tibbs Cortes *et al*., 2021). Development of statistical techniques to examine gene-gene interactions struggle with the scale of data presented by sequencing-based GWAS. These weaknesses of statistical GWAS may be responsible for the increasingly acknowledged “missing heritability” problem, wherein a significant proportion of the genetic basis of heritable traits remains unknown (Manolio *et al*., 2009). While continued efforts to improve and refine these statistical GWAS approaches are warranted, it is reasonable to consider entirely alternative approaches to this complex challenge.

Lately, there has been development of both supervised and unsupervised artificial intelligence/machine learning (AI/ML) approaches for performing GWAS using neural networks; these tools likewise utilize discrete variant-to-phenotype relationships as input data and strive to identify meaningful associations (Yoosefzadeh-Najafabadi *et al*., 2022), albeit with fewer algorithmic restrictions and a corresponding “freedom” to find the best approach within the limits of the architecture employed (Sun *et al*., 2020). Neural networks are a type of “deep learning” algorithm that have many applications in the AI/ML space, for instance in image recognition of “computer vision” (CV). CV algorithms are used in problems of identification, when researchers need to discriminate among different classes of entities (i.e., images, shapes, text) from a desired sample group (Al-Saffar *et al*., 2017, Chen *et al*., 2021). At their core, image classification algorithms require training data, in terms of images with correct class labels, which are used to learn the relationship between input pixels and image class. There are numerous algorithms available for that learning process (Lu & Weng, 2007), with convolutional neural networks (CNNs) being one of the most commonly used (Rawat & Wang, 2017). Researchers and developers have created image classification algorithms that are pre-trained on millions of labeled images, which can speed the process of developing classification schemes and applying them to new problems (He *et al*., 2016). Furthermore, there are a number of existing “post-processing” algorithms that can be used to determine the influence of individual pixels to an overall classification prediction; these methods, which include Local Interpretable Model-Agnostic Explanations (LIME), Shapely Values, layer relevance propagation, and integrated gradients (IGs), aid in interpreting image classification model activities (Sundararajan *et al*., 2017). Overall, CV based on ML is a well-established computational tool for classifying images to solve a variety of challenges.

Recently, CV has been applied to problems in genomics, ranging from variant identification (Poplin *et al*., 2018) to gene expression analysis (Wang *et al*., 2021). A central problem in genomics is understanding the relationship between the genome and resultant phenotypes. With accurate genotype-to-phenotype maps, researchers could reasonably predict the observed traits of an individual organism simply through its genome (though environmental variation can play a major role in phenotype as well). While deep learning and CV have been applied to phenomics – the automated identification and discrimination of distinct phenotypes (Houle *et al*., 2010) – CV has not historically been applied the genome itself. CV, however, may be well-suited to learning complex associations between genotype and phenotype (Ubbens & Stavness, 2017, Lürig *et al*., 2021), provided that visual representations of the genome can be fed into a CV algorithm (Muneeb *et al*., 2022). In this way, CV may provide advantages over traditional GWAS, which tend to be computationally burdensome and have limited value in determining causal associations between genes and phenotypes (Witte, 2010, Korte & Farlow, 2013, Tam *et al*., 2019).

Typical AI/ML approaches for genome variation analysis work on a matrix of shared single nucleotide polymorphisms (SNPs) across a population, while not taking positional information into account; different variants located in the same gene or even at the same position are inherently treated as separate data categories (Ma *et al*., 2018) unless alternate algorithms are employed. For AI/ML CV tools, this relational information is critical. A CV algorithm, for instance a CNN, will typically consider the patterns formed by elements (pixels) when solving a classification challenge, and use patterns in the elements to discriminate between different output classes (e.g., objects). Consideration of these patterns, as well as their relation to each other in a physical genome, provides a potential advantage for CV over traditional GWAS when used to understand genomic causes of phenotypic changes; CV can plausibly use more information than an existing GWAS pipeline, enabling the statistical modeling of more complex interactions. CV, then, may represent a powerful but untapped resource for genotype-to-phenotype prediction and association detection, albeit one that has not yet been adapted to perform such tasks (e.g., one notable weakness in this context is that current CV tools, by design, assign low prioritization to pixel location in the image (LeCun *et al*., 1995), a notable disadvantage in the GWAS context).

This concept was tested by translating genomic variation into images, then using a state-of-the-art CNN-based CV approach to learn the relationship between images and resulting phenotypic variation. Existing image-training databases and algorithms were used for model development and a proof-of-concept genotype-to-phenotype prediction pipeline was developed in an agriculturally important species (maize).

## MATERIALS AND METHODS

### Data Collection

To develop a proof-of-concept CV pipeline to translate genomic variation to phenotype predictions, an existing dataset was required with: 1) available information on genomic variation; 2) associated phenotype information for different variants, strains, or genotypes; and 3) an existing body of GWAS results that could be used to validate and compare the analysis.

Many GWAS studies in maize (*Zea mays*) have been reported to date (Xiao *et al*., 2017); however, the majority of these studies have, for historical and economic reasons, been restricted to microarrays or SNP panels. SNP panels are cost-effective sequencing approaches that are restricted to ∼50K SNPs and include only exonic and intronic coding regions, not intergenic (Xu *et al*., 2017). Intergenic regions can include key elements involved in modulation of gene expression, equally important for phenotype determination (Tonnessen *et al*., 2021). Coincident with the exponential decreases in sequencing costs, recent years have seen the rise of full-genome sequence-based GWAS studies in maize. Recently, Mural et al. gathered all available sets (15 in total) of maize genome resequencing data associated with trait values, over 1,000 samples in total, and made a novel effort to make them fully interoperable by calling variants for each against the Maize B73 version 4 reference genome (Mural *et al*., 2022). Mural et al. made the variants for all samples available as a single variant calling format (VCF) table, an ideal GWAS resource for development and testing of the techniques employed in this study.

### Genomes to Image Conversion

The conceptual approach to decomposing genomic variation into images involves the use of *k*-mers, which differs from the SNP-based image approach of Muneeb et al. (2022). For the maize data, the large VCF file published by Mural et al. (2022) was used. Each VCF file was split into windows (N = 500,000 lines), which were then decomposed into smaller sub-windows (N = 5,000 lines) for counting *k*-mers. Within each sub-window, the reference alleles and variant alleles (if present) encoded in the VCF file were concatenated, and these sequences were used to count *k-*mers within each reference and variant sequence segment. After *k-*mers were calculated within each sub-window, all pairwise Pearson correlation scores were calculated among sub-windows to generate a correlation matrix for each larger window and this matrix was saved as an image for use as a model input. Each VCF file contained roughly 17 million lines, which were split into windows of 500,000 lines. Within each 500,000-line window (35 windows total), sub-windows of 5,000 lines (100 sub-windows total) were used to concatenate reference and variant alleles (if present) and count *k-*mers of length *k* (3, 5, or 7) within each 5,000 bp concatenated reference and variant sequence. Since there were 100 sub-windows in each larger window, all pairwise correlations were performed between each length 100 vector of *k-*mer counts to generate a 100 by 100 matrix, which was written to file as a grayscale image. Thus, a total of 35 images were generated for each genotype comprised of 100 by 100 pixels with the intensity of pixels representing the correlation between *k-*mers detected in each sub-window. Example images generated by this process are shown in Figure 1 and Figure 2.

**Figure 1.**
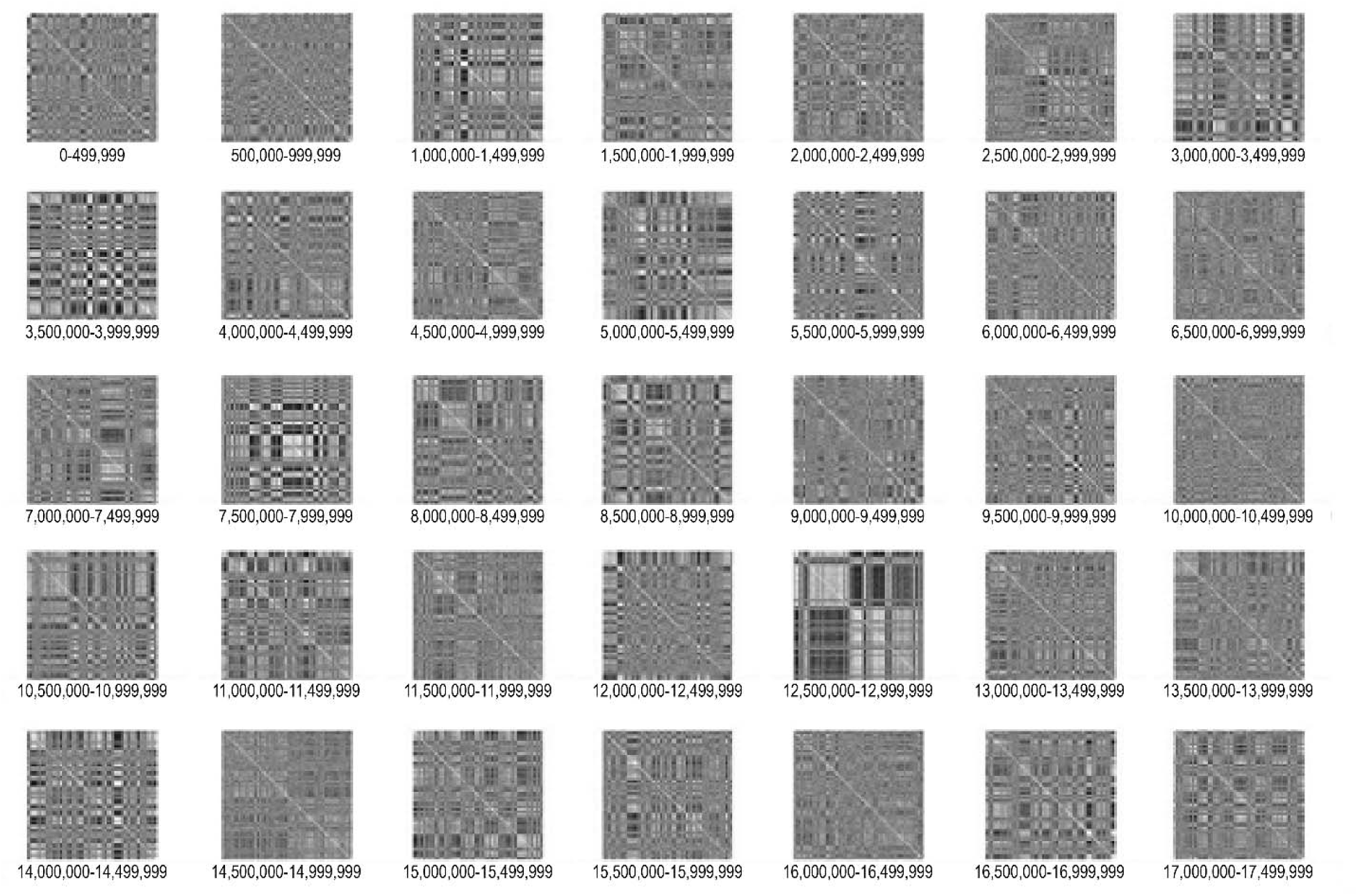
Example of corn k-mer spectral images for a single genotype used as input for training and testing of the ACED CNN algorithm. The image is derived from single nucleotide polymorphism data for corn genotype A385. Each pixel represents Pearson correlations between tetranucleotide k-mers (k = 4) detected within a window of 5,000 variant basepair (bp) segment between corn variant genotype A385 and the reference B73 genome. Pearson correlations were converted to values between 0 and 255. Smaller values are black and larger values are white in the image and are associated with the original Pearson correlation scores. Each image represents a portion of the corn genome, referenced by numbers below each image.

**Figure 2.**
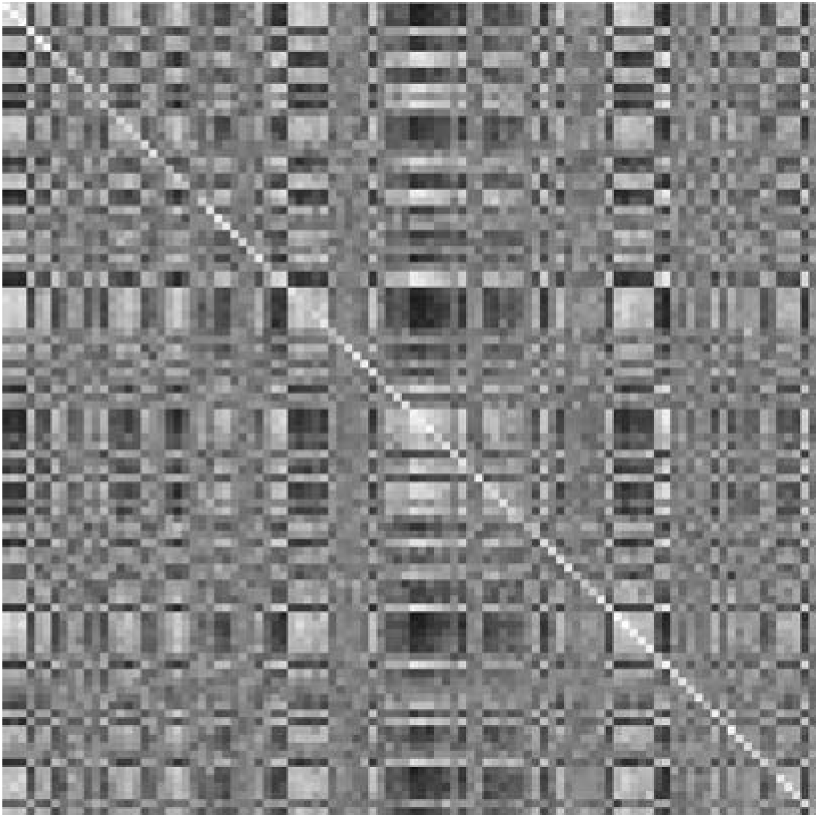
Expanded example of one of the corn k-mer spectral images for genotype A385, which covers a 500,000 variant bp window of genetic variation within the corn genome.

### Selection of Genotypes Representing Heterotic Groups

In the original dataset, Mural et al. provided the heterotic group assignment for each genotype, including Non-Stiff Stalk, Stiff Stalk, Tropical, Sweet Corn, Popcorn, Iodent, and Flint. For assessing the technique’s ability to ingest the genome *k-*mer spectral images and properly perceive at least some degree of genomic variation patterns, a classification model was trained on the three most populous groups, Non-Stiff Stalk (n=300), Stiff Stalk (n=221), and Tropical (n=116).

### Sub-Selection of Genotype-to-Trait Data for Phenotype Classification

The second phase of model construction was aimed at classifying trait values within a group of genotypes representing a minimally structured population. The Non-Stiff Stalk heterologous group was chosen because it was the largest among those available (n=300). Three traits with high sample count (n) and different degrees of heritability (H) were chosen for model training under the hypothesis that, if the model were keying in on real genetic trait drivers, heritability would correspond to model performance. Trait H values were derived from Mural et al. Ear height, plant height, and tassel length were chosen as the traits with high (n=225, H=0.73), medium (n=130, H=0.53), and low (n=247, H=0.25) heritability, respectively. Trait values were subjected to *k*-means clustering with *k* = 3 clusters to generate labels for three value levels – low, medium, and high.

### Model Architecture and Training

EfficientNet was utilized as the main backbone of the model training system. Compared to other popular networks such as ResNet and AlexNet, EfficientNet shows the best performance while also minimizing the number of parameters and floating-point operations (flops) needed (Tan & Le, 2019). EfficientNet has been trained on the ImageNet-1K challenge (Russakovsky *et al*., 2015), which consists of approximately 1 million real-world images that have been classified into 1,000 distinct classes. Different complexity levels of EfficientNet have also been developed, starting from B0 (which performs on par with ResNet-50), up to B7. The model can also allow images of any size as input. However, these images are RGB (i.e., 3-channel data). The input had 36 channels, which is much larger than the design specifications for EfficientNet. To accommodate the extra channels, the first 3 channels of the image were extracted and fed through an EfficientNet-B7 model. The B7 model’s weights were frozen to prevent further training. The remaining 33 channels were fed through a separate EfficientNet-B0 model. The first 2D convolutional block was modified to accept the increased number of channels. This B0 model was *not* frozen so that it finetuned during training. Both 1,000-dimensional outputs were then concatenated and reduced to three classes to match the output size of the two tasks; this minimal linear block was also allowed to train. Figure 3 shows the final architecture of the model, called ParallelNet. Training and validation were performed using a desktop workstation equipped with an Intel Xeon Gold 5220 CPU and NVIDIA Quadro P1000 and P5000 graphics processing units (GPUs).

**Figure 3.**
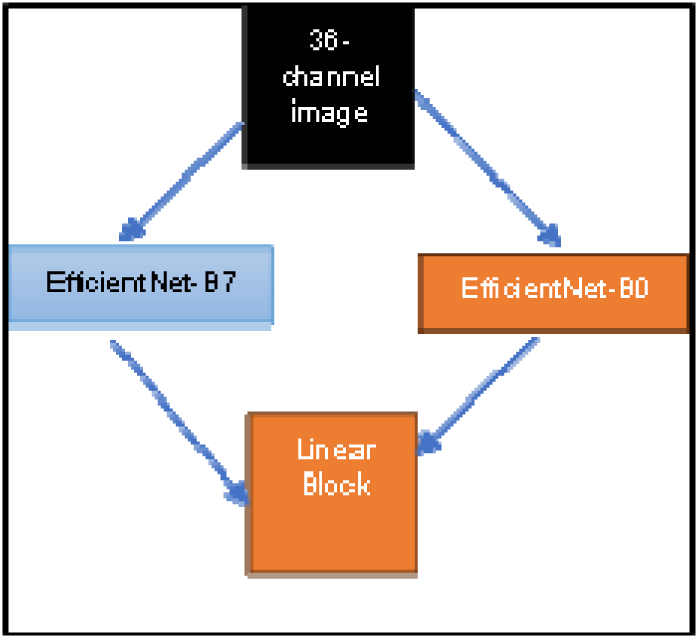
Schematic of the model architecture, “ParallelNet,” which was applied for training the genome image classification model.

During training, *k*-fold cross validation was used, applying *k* = 5 groups, which allowed for the creation of a full set of validation metrics while avoiding data leakage. In each iteration, 80% of the data were used for training, and the remaining 20% used as validation. During the final step, all validation metrics were aggregated. Models were trained using the Adam optimizer with a learning rate of 0.001 (Zhang, 2018).

### Feature Attribution Extraction

After training and validation were completed, models were examined to determine which image pixels, and thus which genome regions, were most informative for classification. To aid in pinpointing the exact regions of the genomes that contributed most toward trait-level predictions, the IG method was employed (Sundararajan *et al*., 2017). This algorithm takes as input the model parameters, a specific trait level target, a full 36-channel input image, and outputs for each pixel in the original image to generate the relative contribution toward the raw trait-level prediction (hereafter attribution score). Negative attribution scores indicate areas of the image that contributed against placement into the target trait class, while positive attribution scores indicate areas that contributed toward placement into the target trait class.

The IG algorithm produced attribution values for each pixel (100 by 100 per image) across each genome-window image (n = 35 per genotype), using input from each of the corn variants with measurements for each of the selected phenotypes. IG was applied to each of the three phenotypic traits and three levels of each trait (low, medium, high). The mean attribution of each pixel (range of genome locations) to image classification was calculated as the sum of the mean values across all rows at each location, resulting in a “sum(mean)” attribution value for each genome region.

### Pathway Mapping and Enrichment

Considering the large size of the corn genome, only genome locations with variant sites across the full dataset were considered in the image generation and model construction, at a pace of 5,000 genomic sites (variant or reference alleles) per pixel, which reduced the input data from ∼300 million basepairs to ∼17 million variant sites. Thus, each pixel did not correlate with a fixed genome region; average total bp covered by each pixel was ∼590,000 bps and genome “pixels” contained 11.3 ± 7.5 genes.

Mapping of maize genes to biological pathways and annotation categories was provided by the Plant Reactome database (PRdb) (Naithani *et al*., 2016, Naithani *et al*., 2019), the Maize Genetics and Genomics Database (MGDB) (Portwood *et al*., 2018), and the Kyoto Encyclopedia of Genes and Genomes (KEGG) (Kanehisa *et al*., 2016). MGDB was generated using the Ensemble Enzyme Prediction Pipeline of the strain B73 *Z. mays* genome. This gene-to-pathways assignment was generated on the latest version of the corn genome, version 5, as opposed to version 4, which was used in the Mural et al. project, and thus presently. Transference of the annotations from the v5 to v4 genomes using a mapping table provided at MGDB generated 5,647 pathway mappings for 2,568 genes. When both the MGDB and PRdb databases are considered, a total of 3,442 genes have at least one pathway mapping, constituting ∼9% of the total genes encoded in the corn genome. Additionally, 14,803 genes were assigned to one or more KEGG pathways by direct genome annotation using the BlastKoala pipeline (Kanehisa *et al*., 2016). Figure 4 provides a schematic of the pathway enrichment workflow.

**Figure 4.**
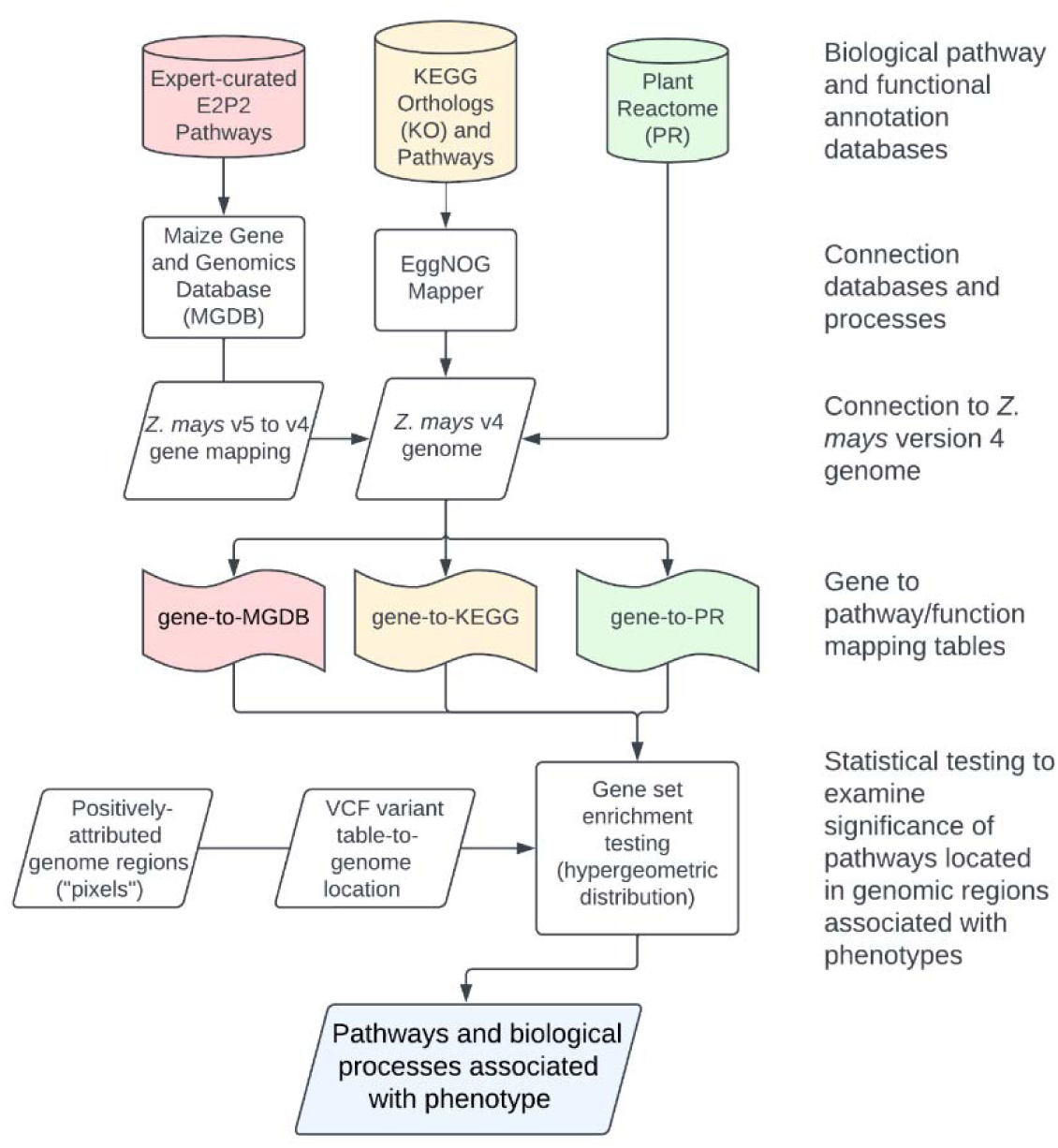
Methodological flowchart detailing the assignment of biological pathways to regions of the genome determined by the corn model to be informative for genotype-to-phenotype classification.

## RESULTS AND DISCUSSION

### Heterotic Group Classification

The heterotic group classifier performed with an overall accuracy of 83.4% for all the validation samples, with similar scores for the recall metric (Figure 5). The largest errors occurred when the model predicted Tropical when the actual group was Non-Stiff Stalk, and when the model predicted Non-Stiff Stalk for certain Stiff Stalk plants. Correspondingly, Mural et al. also reported some overlap in in the genetic relatedness between the three groups when subjected to all-SNP multidimensional analyses (Mural *et al*., 2022).

**Figure 5.**
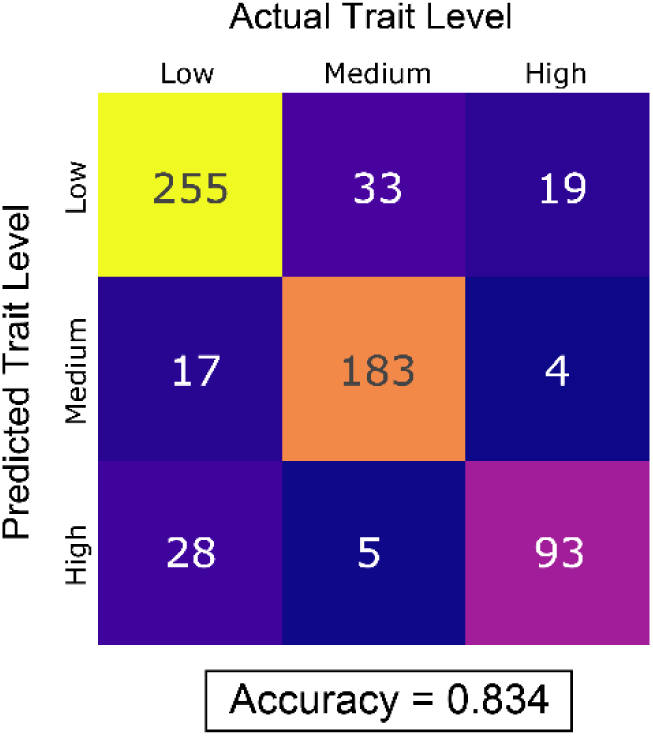
Validation matrix for the heterotic group classification model.

### Trait Group Classification

#### Distributions of Trait Values

For evaluating the approach’s ability to predict phenotype, three traits were chosen from the largest heterotic group, ear height (EarHeight_L), plant height (PlantHeight_M), and tassel length (TasselLength_J), as representatives with high, moderate, and low heritability, respectively. The distribution of these trait values exhibited characteristics of a normal distribution (Figure 6). The long right tail of TasselLength_J is the only exception to this trend.

**Figure 6.**
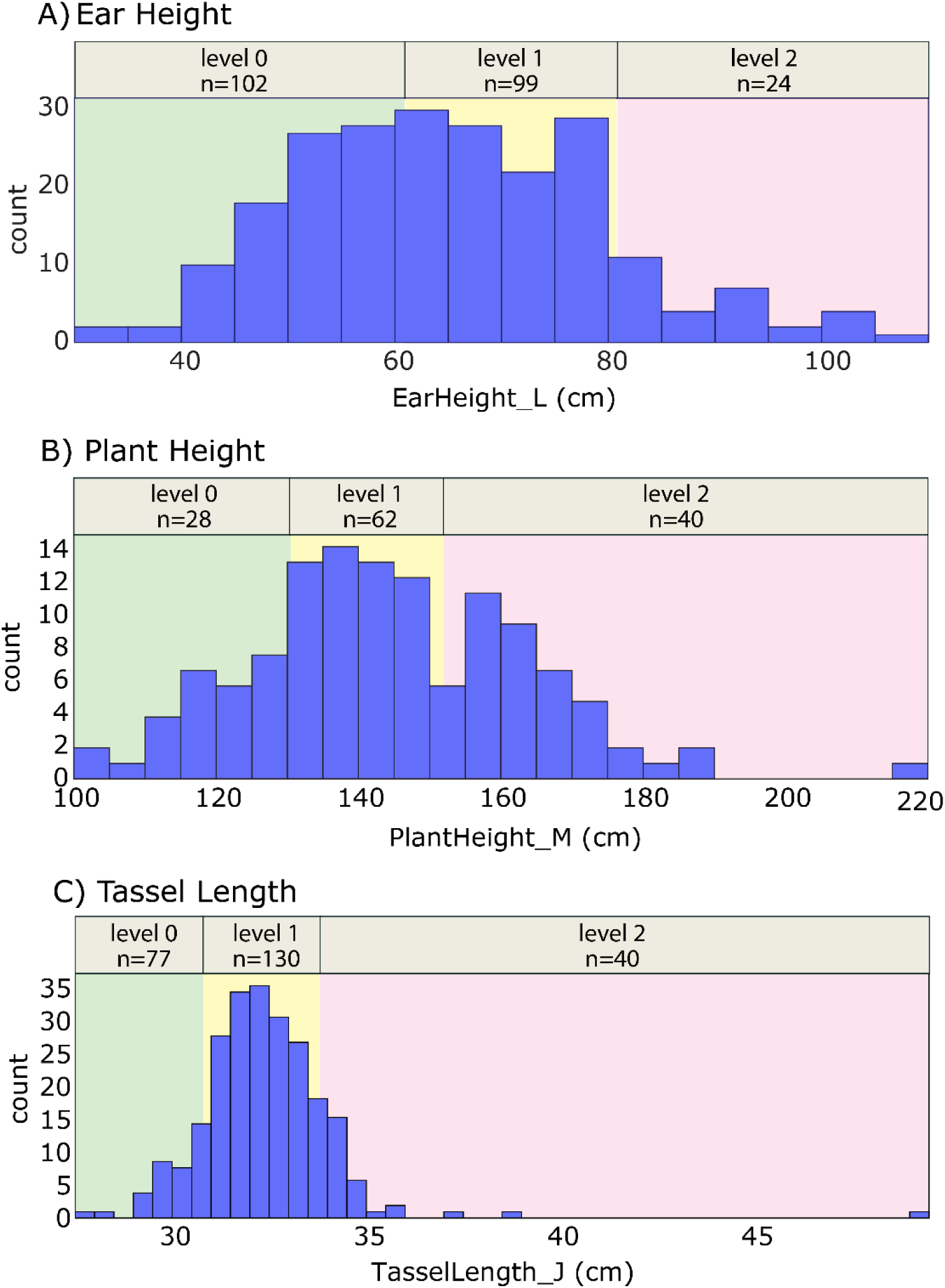
Distributions and k-means clustering results for the three selected traits in the Non-Stiff Stalk heterologous group. A value of k=3 for the k-means clustering operation. The letter “n” indicates the total number of individual genotypes placed into each cluster.

#### 2. Validation Results

The trait-level classification task did not perform as well as for heterotic groups (Figure 7). The main errors occurred when predicting between the two most common trait levels in for each trait. The ear height model displayed the best ability to differentiate between low and medium trait levels, while plant height and tassel length showed only moderate success across all differentiations, consistent with the lower heritability of these traits. Also, this result is partially due to the relatively low number of trait levels in the minority class, which incentivizes the model away from predicting too many genotypes with that trait level, even with the included loss weighting. Additionally, the distribution characteristics of the trait levels exhibited those of a normal distribution. By its nature, the true trait level of most samples will be classified as medium, with lower sample numbers for both low and high. The direction of the skew will determine which class has the lowest representation, and to a lesser extent, the initial center values set by the *k*-means clustering algorithm. The placement of continuous trait values into distinct clusters led to a significant loss of real biological data, but was required for this initial prototype approach. Regardless, the fact that each model showed above-random success (with random success = 0.33) shows promise for the approach.

**Figure 7.**
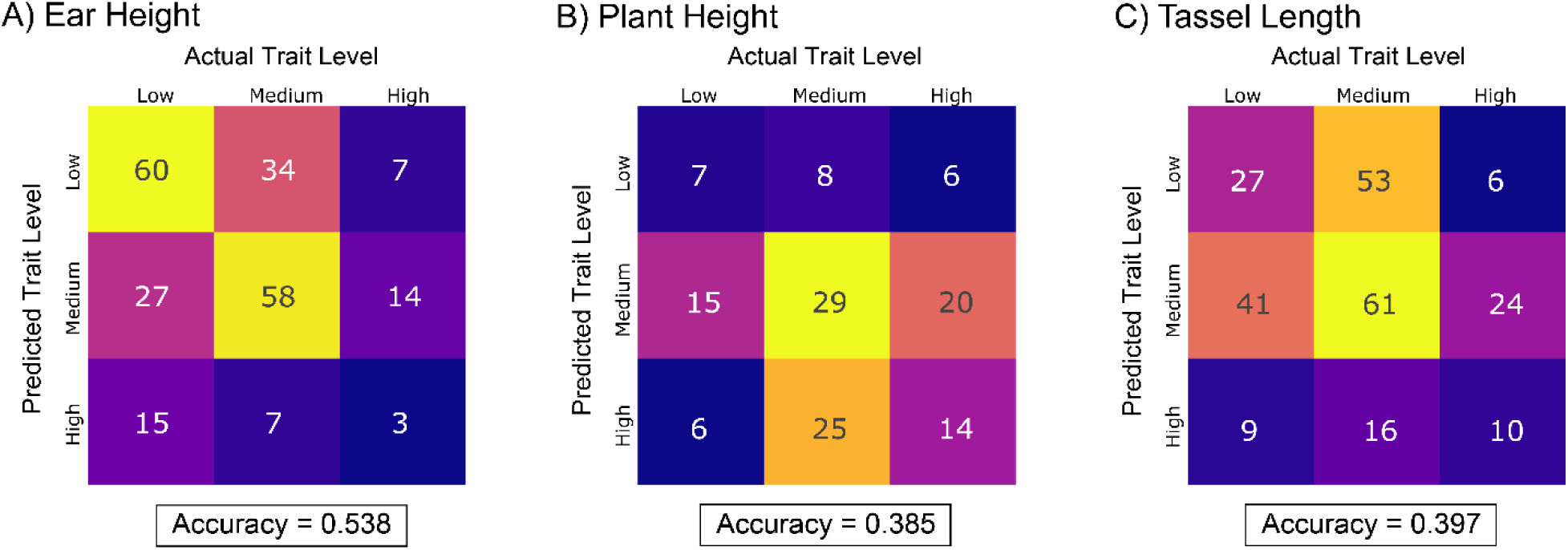
Validation results for the three Non-Stiff Stalk heterotic group trait phenotype classifiers.

#### 3. Extraction of Attribution Scores

Given the relatively high predictive performance of the Non-Stiff Stalk ear height classifier, IG was used to determine which areas of the maize genome contributed to trait value group predictions. Figure 8 shows an example genome *k-*mer spectral image and the resulting attribution image for one genotype (strain WD) for classification into the low ear height category. Positive attribution scores indicate spectral image regions that contributed to a decision to place the genome into the category under examination, while a negative score indicates influence to not place the genome into that category. Three (3) distinct attribution tables were generated for each trait value category (low, medium, high).

**Figure 8.**
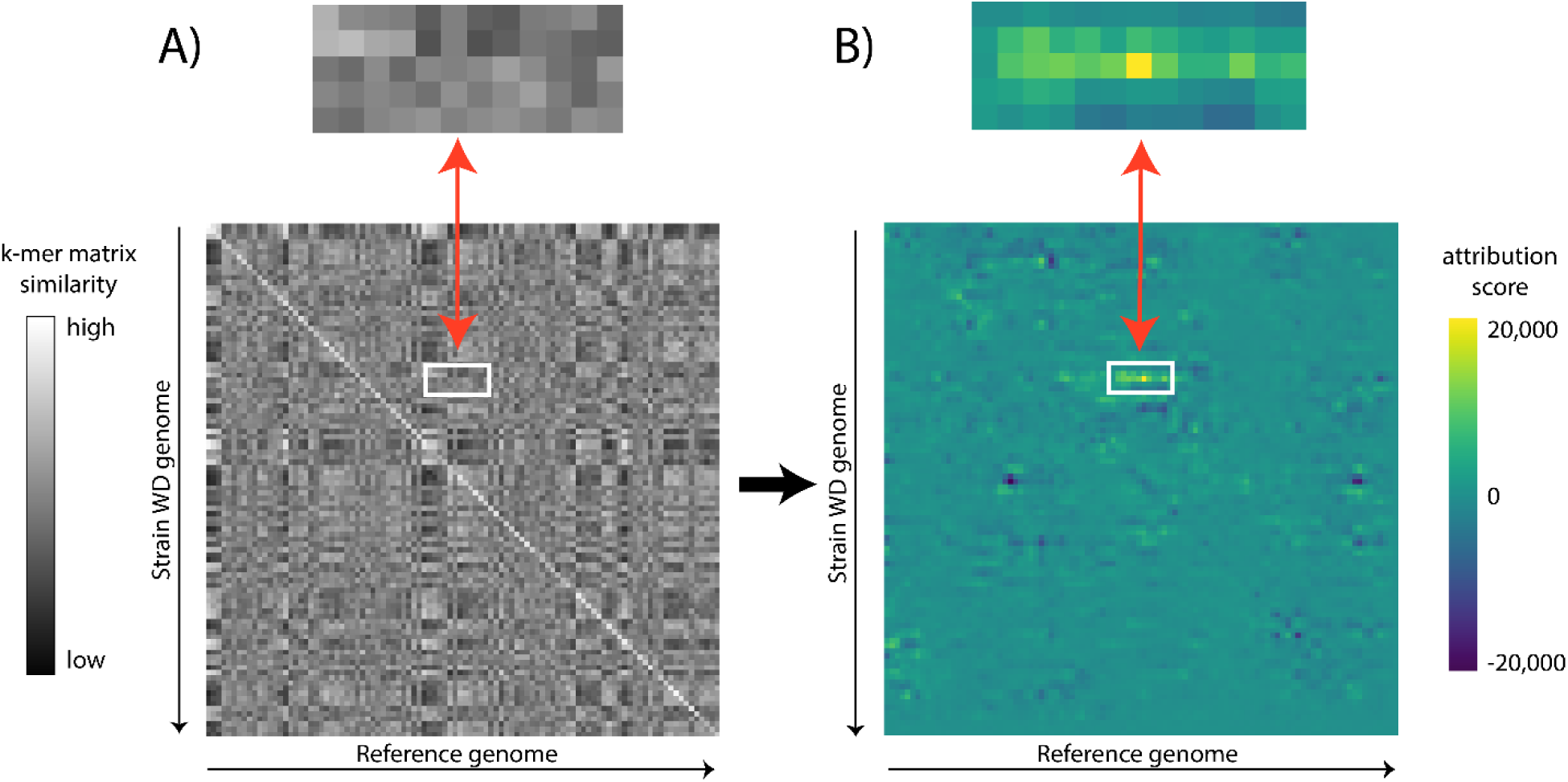
Example depicting the generation of A) the first of 35 k-mer spectral images for the genome of Non-Stiff Stalk strain WD and B) heat plot representation of the attribution scores for the model’s decision to classify as low ear height by the model. The boxes above each image show an expanded sub-region of the image containing the region which was assigned the highest attribution score.

To generate metrics reflecting which genome regions tended to contribute most strongly to the model’s classification decisions across all strains, attribution scores were aggregated across genotypes. While the inverse x-y intercept represents the precise comparisons between corresponding regions in the sample and reference genomes, changes in the spectral image at those points of intersection also lead to cascading changes across the remainder of the position-to-position comparisons, which serves to amplify the signal, but may also lead to the classifier learning to focus on parts of the image away from this precise intersection, a phenomenon that can be seen in the attribution image pictured in Figure 8. To account for this trend, the mean value across each genome position was taken as a final scoring for each position within each genome. To aggregate this scoring across all genotype attributions for each trait category prediction, these values were summed and examined as a whole for positions with outlier values (defined here as two standard deviations above or below the mean regional value), which provides the ability to determine which genome regions have higher or lower aggregated attribution scores than a typical region. These outliers, when traced back to specific genome locations, represent the approach to flagging impactful genome regions that are consistently associated with certain output classes.

#### 4. Biological Pathways in Meaningful Genome Regions for Ear Height

Because it was the best performing model and represented the most heritable trait, the ear height predictions were targeted for extraction of meaningful genome region attributions; 4.5-5.7% of the genome was flagged as meaningful by this method, reflecting the fact that a two-standard-deviation approach was applied to flag outliers (Figure 9).

**Figure 9.**
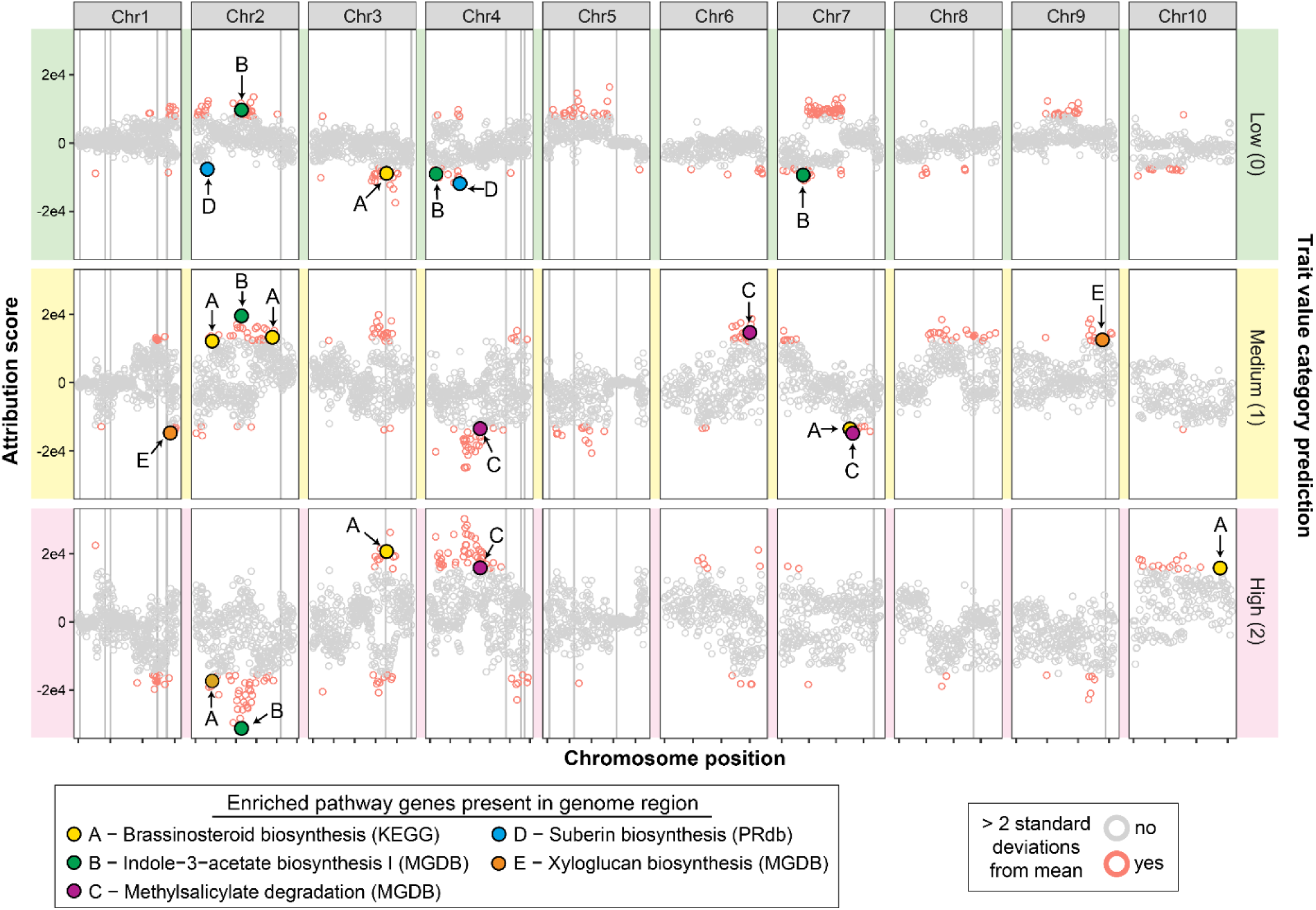
Aggregate attribution values for each genome region derived from the EarHeight_L model. Regional scores for each of the three categorical trait value predictions are presented separately in the top, middle, and bottom panels as indicated. Each vertical panel represents 1 of the 10 main Z. mays chromosomes as named at the top. The 17 vertical lines indicate regions identified by Mural et al. as informative in their GWAS study of the same genotypes and trait data. Regions flagged as potentially informative are differentially colored according to the bottom right legend. Informative regions containing genes contained in statistically enriched pathways are flagged with colored circles and labeled according to the legend at the bottom left.

This proportion of the genome contained 1,510-2,037 genes. To assess the biological relevance of this genome sub-region, pathway enrichment analyses were conducted (considering only pathways containing at least two genes). Only one pathway, KEGG Ribosome, passed a multiple-testing-adjusted p-value cutoff of 0.05 for enrichment, albeit with a small enrichment factor (EF) of 1.9. An additional 66 pathways from the three systems applied (KEGG, MGDB, PRdb) were enriched in at least one of the three trait level model attributions with an unadjusted p-value of 0.05 (Table 1). The biological context of most of these pathways represented core metabolic functions, such as amino acid, cofactor, or sugar biosynthesis, making interpretation challenging. However, there were a handful of highly enriched pathways with clear associations with growth and height. EF values were high for pathways for metabolism of hormones involved in plant growth; brassinosteroid biosynthesis (Hu *et al*., 2017) (EF=3.8-4.7, KEGG pathway 00905), indole-3-acetic acid biosynthesis I and II (Zhao, 2010) (EF=3.6-5.6, MGDB), and methylsalicylate degradation (Hu *et al*., 2017) (EF=3.9, MGDB). Pathways for the structural polymer xyloglucan (Hu *et al*., 2017) (EF=5.7, MGDB) and apoplastic water-conservation polymer suberin (Ranathunge *et al*., 2011, Woolfson *et al*., 2022) (EF=4.9, PRdb) were also highly enriched. The full list of enriched pathways is provided in Supplemental Table S1.

**Table 1.**
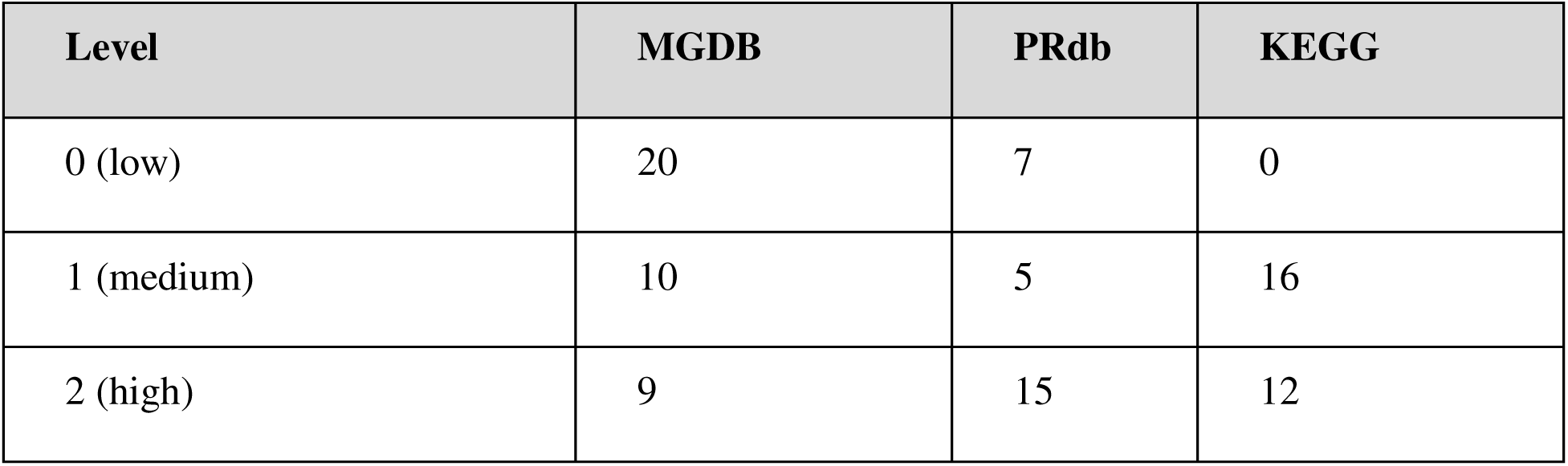
Total number of enriched pathways in the meaningful genome attributions utilized by the three trait level classifiers. Only pathways with at least two genes and with an unadjusted p-value < 0.05 were considered.

The biological plausibility of the enriched pathways in the most informative genome regions may suggest that querying the trait category classifier offers a novel technical approach to GWAS, although conclusions should be tempered by the fact that no efforts to control for linkage disequilibrium were applied. Conversely, the informative regions as a whole did not coincide well with genome regions identified the formal GWAS conducted by Mural et al. as determined by application of the FarmCPU algorithm (Liu *et al*., 2016); only 3 meaningful regions as determined by model attribution scores coincided with the 17 regions identified by Mural et al. (Figure 9).

### Outcomes and Prospects for CV-Aided GWAS

The CV model was reliably able to separate phenotypes by genetic lineage (i.e., heterotic group), but that performance predicting individual phenotypes was mixed. CV-informed GWAS offers several potential advantages over statistical GWAS, and can be improved by: 1) increasing the quality and availability of relevant input data; 2) training the models on a greater variety of phenotypes (including true damage phenotypes) with higher resolution genome-encoding; and 3) developing purpose-built CV algorithms that are trained on genome images.

AI/ML technologies are rapidly supplanting complex analytic tasks in all areas, especially where data streams are increasingly vast and statistical analyses overwhelming, computationally and intellectually. GWAS is an ideal candidate for development of AI/ML solutions; genomes are large with complex layers of interactions that determine phenotype, the variety of genotypes is essentially infinite and ever-changing, and the cost of genome sequencing is decreasing at an exponential rate. Statistical GWAS has succeeded in identifying critical drivers of many traits; however, it is widely acknowledged that the majority of genomic determinants of trait heritability remains unknown and as-yet unsolvable. The modest success observed was in the present study and should be interpreted in the light that: 1) the off-the-shelf CV tools that were applied were not well-suited to the task; 2) the best way to parse the attribution scores resulting from model result querying has not been identified; and 3) sample counts and image sizes were quite smaller than ideal for training such models. Despite these limitations, the prototype model was able to predict a complex phenotype at an above-random success rate and map those regions back to biologically plausible biological pathways.

Well-developed CV-aided GWAS technologies offer the prospect of rapid prediction of the phenotype of novel genomes, detection of aberrant genomes (products of hybridization or engineering), and gene-to-phenotype associations in a manner that is easier to operate than current GWAS tools and takes advantage of the rapid development of GPU-based hyperthreaded computational systems. Further, because deep learning models have the freedom and creativity to solve classification challenges by simultaneous application of a large variety of logical and non-linear mathematical approaches that take not only major signals, but also subtle covariance patterns into account, they may offer hope for solving at least a portion of the missing heritability problem.

### Limitations and Areas for Improvement

#### Sample Size

The sample size of the current dataset posed a hurdle in the learning ability of the network. While transfer learning was performed by using existing pre-trained architectures, these models were trained on massive datasets. The popular ImageNet dataset consists of over 1 million images from which the model will train (Russakovsky *et al*., 2015), while the dataset numbers fewer than 400, which likely led to increased difficulty in the model’s ability to generalize to unseen genotypes. Additional data comprised of genome variants and resulting phenotypes would benefit model training and subsequent predictions.

Additionally, the model was trained on “normal” maize phenotypes (see Figure 6). That is, the separation of phenotypes into bins of “low,” “medium,” and “high” trait values was somewhat arbitrary, and may be challenging for genotype-to-phenotype prediction approaches. Other methods of generating paired genotype-phenotype variants may provide better data for testing the utility of this approach. In rice, for instance, researchers have intentionally irradiated plants to generate thousands of mutant lines (Li *et al*., 2017). A limited trial of this methodology was conducted using these mutants, but found insufficient available phenotypic data for the CV approach; additional phenotypes linked to intentionally created mutant lines would provide a useful test case for this concept.

#### Image Dimensions

Popular modern CV image classification tools are trained on images with dimensions even smaller than that provided by typical digital photography. For example, the original EfficientNet was trained on the ImageNet dataset, which consists of 224 by 224 pixel images (Russakovsky *et al*., 2015). Conversely, even the smallest genome represents a vastly larger challenge. The maize reference genome is comprised of over 2 billion bps, which, if translated directly into a 2D mapping image, would include ∼14 orders of magnitude more information than an ImageNet picture, no doubt far exceeding capacities of current tools and computational hardware. To make image dimensions manageable for this construction of this prototype, this study focused only on sites with at least one variant across the dataset and binned variant sites at a rate of 5,000 per bin, which allowed generation of a 35-channel, 100 by 100 image per genotype, which is still approximately one order of magnitude larger than a typical 3-channel 224 by 224 image. This level of data compression represented a significant compromise in biological granularity, but even at this level, the larger image size represented a significant computational burden that necessitated development of a refined architecture (ParallelNet).

Inputs to the trained models were 35-channel images, each channel 100 by 100 pixel. However, the original EfficientNet was trained on the ImageNet dataset, which consists of 224 by 224 pixel images. Therefore, the model’s weights were trained to recognize features at this resolution. Although the network accepts images of any resolution, providing a smaller resolution image can cause the model to have difficulty in differentiating features in the image. One direct solution is to develop a methodology to use images that would be much closer in size to the original 224 by 224 pixel image.

#### Model Architecture

A fundamental limitation of this approach is the use of existing CV architectures for model development and training. The EfficientNet B0 and B7 architecture, for instance, has been built for common CV needs, such as locating people, animals, or vehicles in complex backgrounds, and recognizing emotional states. It is pre-trained on the benchmark ImageNet dataset. However, ImageNet consists of real-world images, in contrast to the manufactured *k*-mer correlation maps fed as the input. It was suspected that a CV architecture built for and trained on the type of genomic images used in the pipeline, validated with known phenotypes, and linked to well-annotated biological pathways would yield better performance than an architecture trained to distinguish between dogs and cats. A beneficial next step in this area would be to develop an architecture and associated training image set that is built specifically for phenotype predictions from images of genomic variation.

There are several other architecture-related limitations that were identified through this work, alongside opportunities for improvement. First, the ParallelNet architecture developed provides a method to handle the large number of channels of the input. Both branches of ParallelNet consist of EfficientNet variants. However, there are many other pre-trained image recognition models. The transformer and attention mechanisms that displayed great performance in the text model space has also been implemented for image recognition models. These models include the Vision Transformer (Alexey Dosovitskiy, 2020) and the Swin Transformer (Liu *et al*., 2021). These models are more complex and require more computing resources and train and validate, but would represent plausible alternatives to improve model predictive performance.

Additionally, an ideal architecture for addressing CV problems related to the genome needs to place value on the absolute position of pixels in an image, as well as relationships among pixels because pixels in an image of a genome have a clear physical relationship to positions (and genes) in the genome. For example, while the absolute position of a steering wheel in an image may not matter to an algorithm attempting to identify a car, the absolute position of an impactful region of the *k*-mer spectral images has a biological basis that must be accounted for in model architecture.

## ACKNOWLEDGMENTS

This study was funded by the Defense Advanced Research Projects Agency (DARPA) Defense Sciences Office (DSO) Program DARPA-PA-22-01-01 Foundational Systems for Food Security (FS2) program under agreement HR0011-23-9-0055. The views, opinions, and/or findings expressed are those of the author(s) and should not be interpreted as representing the official views or policies of the Department of Defense or the U.S. Government

## SUPPLEMENTAL MATERIAL

Supplemental Table S1. All enriched pathways (unadjusted p-value < 0.05) encoded in genome regions corresponding to meaningful image pixels as determined by the Ear Height classification model. “description” is the name of the pathway in the given “database”. “trait_level” indicates whether the genome region corresponds to low (0), medium (1), or high (2) trait level classification. “subset_ocurrences” indicates the number of genes in the pathway were in the impactful genome region and “subset_total” is the total number of genes with mappings to the indicated database are encoded in the impactful region. “genome_occurrences” is the total number of genes mapped to the pathway in the entire genome, and “genome_total” is the total number of database-mappings in the entire genome. “enrichment_factor” is the degree of enrichment of the pathway in the impactful genome region as determined by hypergeometric testing, with the associated “p_value” provided in the adjacent column. “p_adjust” is the result of adjusting the p-value using a multiple testing correction.

## DATA AVAILABILITY

All data and models are available from the authors upon request.

## AUTHOR CONTRIBUTIONS

The authors acknowledge contributions to the paper are as follows: study conception and design: SAH, MM, CD, RWM; data collection: SAH, RWM; analysis and interpretation of results: SAH, EA, MM, CD, RWM; draft manuscript preparation: SAH, RWM, CD. All authors reviewed the results and approved the final version of the manuscript.

## Abbreviations

AI: artificial intelligence
CNN: convolutional neural network
CV: computer vision
EF: enrichment factor
GPU: graphics processing unit
GWAS: genome-wide association studies
IG: integrated gradient
KEGG: kyoto encyclopedia of genes and genomes
MGDB: maize genetics and genomics database
ML: machine learning
PRdb: plant reactome database
VCF: variant call format

